# Hippocampal neurogenesis promotes preference for future rewards

**DOI:** 10.1101/399261

**Authors:** Désirée R. Seib, Delane Espinueva, Oren Princz-Lebel, Erin Chahley, Stan B. Floresco, Jason S. Snyder

## Abstract

Adult hippocampal neurogenesis is implicated in a number of disorders where reward processes are disrupted but whether new neurons regulate specific reward behaviors remains unknown. We find that blocking neurogenesis in rats reduces activation of the ventral dentate gyrus and causes a profound aversion for delayed rewards. Delay-based decision-making restructured dendrites and spines in adult-born neurons, consistent with activity-dependent neuronal recruitment. These findings identify a novel role for neurogenesis in decisions about future rewards, which is compromised in disorders where short-sighted gains are preferred at the expense of long-term health.

## MAIN TEXT

Adult neurogenesis in the dentate gyrus (DG) subregion of the hippocampus has been implicated in psychiatric disorders such as depression^1^ but it remains unclear exactly which behavioral processes depend on new neurons, due to the complex and heterogeneous nature of the disorder. For example, one of the core symptoms of depression is anhedonia, often defined as diminished pleasure or interest in pleasurable activities^2^. However, many behavioral processes are subsumed under this broad definition^3^. Relative to controls, depressed patients place less value on future rewards, which may shift behaviors away from optimal, but delayed, outcomes^4^. Altered prospective behaviors may result from hippocampal deficits. Indeed, hippocampal activity patterns reflect future-oriented behaviors in both rodents^5-7^ and humans^8,9^, amnesics have an impoverished imagination of possible future events^10^ and hippocampal damage biases humans^11,12^ and animals^13-15^ towards immediately-available rewards, at the expense of larger, but delayed, rewards. Whether adult neurogenesis contributes to choices about future rewards is unknown. However, blocking neurogenesis in mice reduces sucrose preference^16^, consistent with a possible role in the anhedonic symptoms of depression.

To test the role of adult neurogenesis in future reward choice, we took advantage of the transgenic GFAP-TK rat model to deplete neurogenesis^17^, and operant tasks that have been optimized for rats. As measured by the immature neuronal marker, DCX, neurogenesis was reduced in valganciclovir-treated (VGCV) TK rats by 94% in the dorsal DG and 77% in the ventral DG, compared to wild type (WT) rats (Fig. 1A-F). Blockade of neurogenesis reduced the overall volume of the dentate granule cell layer, consistent with DG atrophy in depressed patients^18,19^, but was without noticeable effect on subventricular zone neurogenesis, likely due to faster recovery in this neurogenic region during the VGCV-free period of operant testing (Supplementary Fig. 1 and Supplementary Fig. 2).

**Figure 1:**
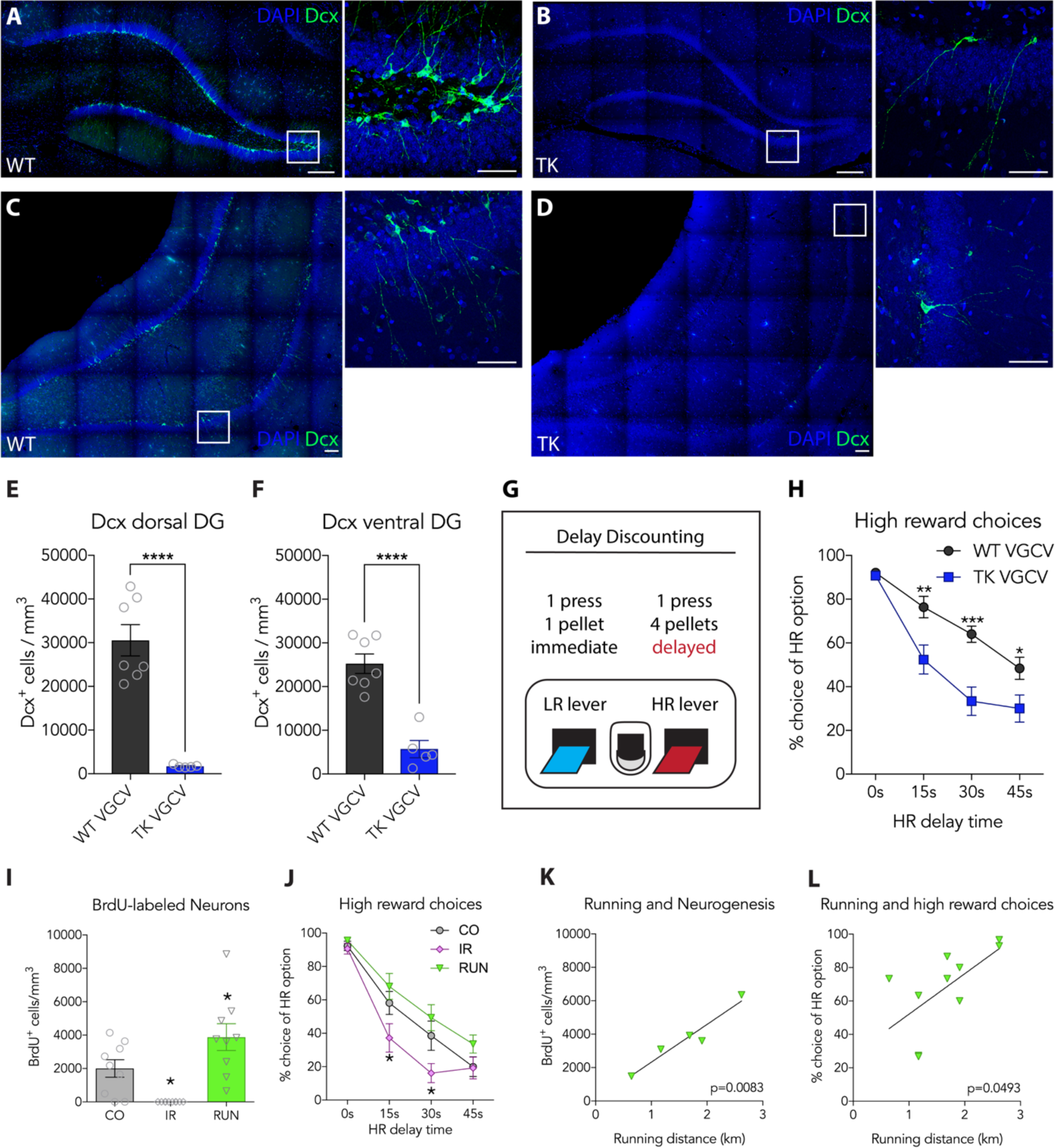
Neurogenesis promotes preference for delayed rewards. Immunostaining for the immature neuron marker Dcx in the dorsal (**A,B**) and the ventral **(C, D)** hippocampus of WT and TK rats. **E, F)** Neurogenesis was reduced in the dorsal and in the ventral DG in VGCV-treated TK rats (T-test, dorsal: WT 30,569 vs. TK 1,738 cells/mm^3^, P<0.0001, n=5-7; ventral: WT 25,255 vs. TK 5,711 cells/mm^3^, P<0.0001, n=5-7). **G**) Overview of the delay discounting operant task. **H)** VGCV-treated TK rats, which lacked adult neurogenesis, preferred the high reward less when a delay was imposed (genotype x time RM ANOVA; effect of genotype: F_1,28_=9.8, P=0.0040; effect of time: F_3,84_=98.6, P<0.0001; interaction: F_3,84_=7.4, P=0.0002; n=14-16; post-hoc Sidak’s test: 15s: WT 76.4 vs. TK 52.41 % HR choice, P=0.0043; 30s: WT 64.02 vs. TK 33.38 % HR choice, P=0.0002; 45s: WT 48.41 vs. TK 30 % HR choice, P=0.0442). **I)** Compared to controls, irradiation depleted and running increased the number of BrdU-labeled cells (ANOVA; effect of group: F_2,25_=13.51, P<0.0001, n=9-10; post-hoc Dunnett’s test: CO 2004 vs. IR 0 cells, P=0.024; CO 2004 vs. RUN 3881 cells, P=0.040). **J)** Irradiation reduced preference for the high reward in the DD task but running did not alter preferences at the level of the entire group (group x time RM ANOVA; effect of group: F_2,27_=3.945, P=0.031; effect of time: F_3,81_=170.7, P<0.0001; interaction: F_6,81_=3.522, P=0.0038; n=10; post-hoc Dunnett’s test: 15s: CO 58.15 vs. IR 37.22 % HR choice, P=0.038; 30s: CO 38.56 vs. IR 16.15 % HR choice, P=0.025). **K)** The number of newborn neurons (born in week 2 of running) was highly correlated with the daily distance run around the time of their birth (weeks 1-3; R^2^=0.9286, P=0.0083, n=5; data points show average values per cage, i.e. pair of rats). **L)** Rats that ran more displayed greater choice of delayed high rewards (R^2^=0.4012, P=0.049, n=10). Graphs display the mean ± standard error for the last 3 test days. Significance indicated between treatment groups and control for I and J. HR, high reward; s, seconds; w, weeks. n indicates number of rats. *, P<0.05; **, P<0.01; ***, P<0.001, ****, P<0.0001.

Delayed rewards typically have less subjective value than immediately-available rewards, a phenomenon often referred to as delay discounting (DD)^20^. To determine if adult neurogenesis regulates the value of delayed rewards, we used a rodent DD paradigm where rats choose between a small reward (1 sugar pellet) that is delivered immediately and a high reward (4 pellets) delivered after a delay^21^ (Fig. 1G). Male rats received 1 session per day for 4 weeks, where each session consisted of 4 blocks of 10 trials. Over the 4 blocks of trials the delay to receive the high reward increased from 0s to 15s, 30s and 45s. During pretraining, when rewards were presented without a delay, both WT and TK rats showed a clear preference (>85%) for the high reward (Supplementary Fig. 2). In the DD task, choice of the high reward decreased with longer delay periods, as expected (Fig. 1H). However, with increasing delays, neurogenesis-deficient TK rats chose the high reward significantly less than WTs. This was not dependent on their motivation to perform the task, since the number of trial omissions was comparable to WT rats (Supplementary Fig. 2). TK rats also displayed normal levels of locomotion and nosepoke behavior, and choice latencies were not different from WTs, indicating that the reduced preference for delayed rewards is not due to non-specific motivational or motor deficits (Supplementary Fig. 2). Furthermore, WT and TK rats that did not receive VGCV were equivalent in all measures, indicating that the delay discounting behavior in TK rats is due to loss of neurogenesis as opposed to off-target effects of the transgenic line (Supplementary Fig. 2).

Adult neurogenesis buffers innate behavioral responses to acute stress^16^ but it is not known if other types of behaviors, such as decision-making, are impacted by acute stress in a neurogenesis-dependent fashion. We therefore subjected rats to 30 min restraint stress prior to DD testing. Performance was similar to unstressed conditions from the previous day, in both VGCV-treated WT and TK rats (Supplementary Fig. 2). These data are consistent with evidence that DD behavior is not impacted by acute stress^21^, and they indicate that neurogenesis depletion does not amplify any latent effects of stress on delayed reward preferences.

To confirm and further explore relationships between neurogenesis and DD behavior, groups of wildtype rats were subjected to irradiation (to inhibit neurogenesis), given access to running wheels (to promote neurogenesis), or left undisturbed in their home cage (Supplementary Fig. 3). Irradiated rats had a complete loss of neurogenesis and they displayed greater DD behavior than intact controls (Fig. 1I,J), similar to what we observed in VGCV-treated TK rats. Irradiated rats were otherwise similar to controls except they were slower to press the lever (Supplementary Fig. 3), which is likely not due to reduced neurogenesis since we did not observe this behavior in VGCV-treated TK rats.

Rats that had access to running wheels ran increasingly greater distances over 4 weeks and had higher levels of neurogenesis than sedentary controls (Fig. 1I and Supplementary Fig. 3). Runners tended to show reduced delay discounting behavior but the group difference was not significant (Fig. 1J). Since there was marked variability in running behavior and neurogenesis, we investigated individual differences and found that rats that ran greater distances also had more neurogenesis and increased preference for the delayed high reward (Fig. 1K,L). Runners were otherwise comparable to controls in terms of omissions, choice latencies, nosepokes and locomotor behavior (Supplementary Fig. 3). These data suggest that, whereas blocking neurogenesis (VGCV-TK and irradiation) can reduce delayed reward preferences, exercise, possibly by promoting neurogenesis, might increase delayed reward preferences.

To determine whether blocking neurogenesis leads to abnormal patterns of activity that could contribute to the reduced preference for delayed rewards we quantified expression of the activity-dependent immediate-early gene Zif268 in the Nucleus Accumbens (NAc; Supplementary Fig. 4), which interacts with the ventral hippocampus during delayed reward choice^22^. There was no difference between VGCV-treated WT and TK rats; both displayed a robust 8-fold increase in Zif268 expression compared to food-restricted cage controls. We next focussed on the long axis of the DG, since the ventral hippocampus has been specifically implicated in delayed reward choice^15^. VGCV-treated WT and TK rats had increased Zif268 activity in the suprapyramidal blade of the dorsal DG after performing the DD task compared to controls (Fig. 2E). In contrast, there was a robust increase in Zif268 expression in the ventral DG of WT rats but activation was severely blunted in TK rats (Fig. 2F). Zif268 expression in the infrapyramidal blade of the DG was not affected by genotype or DD training (Supplementary Fig.4).

**Figure 2:**
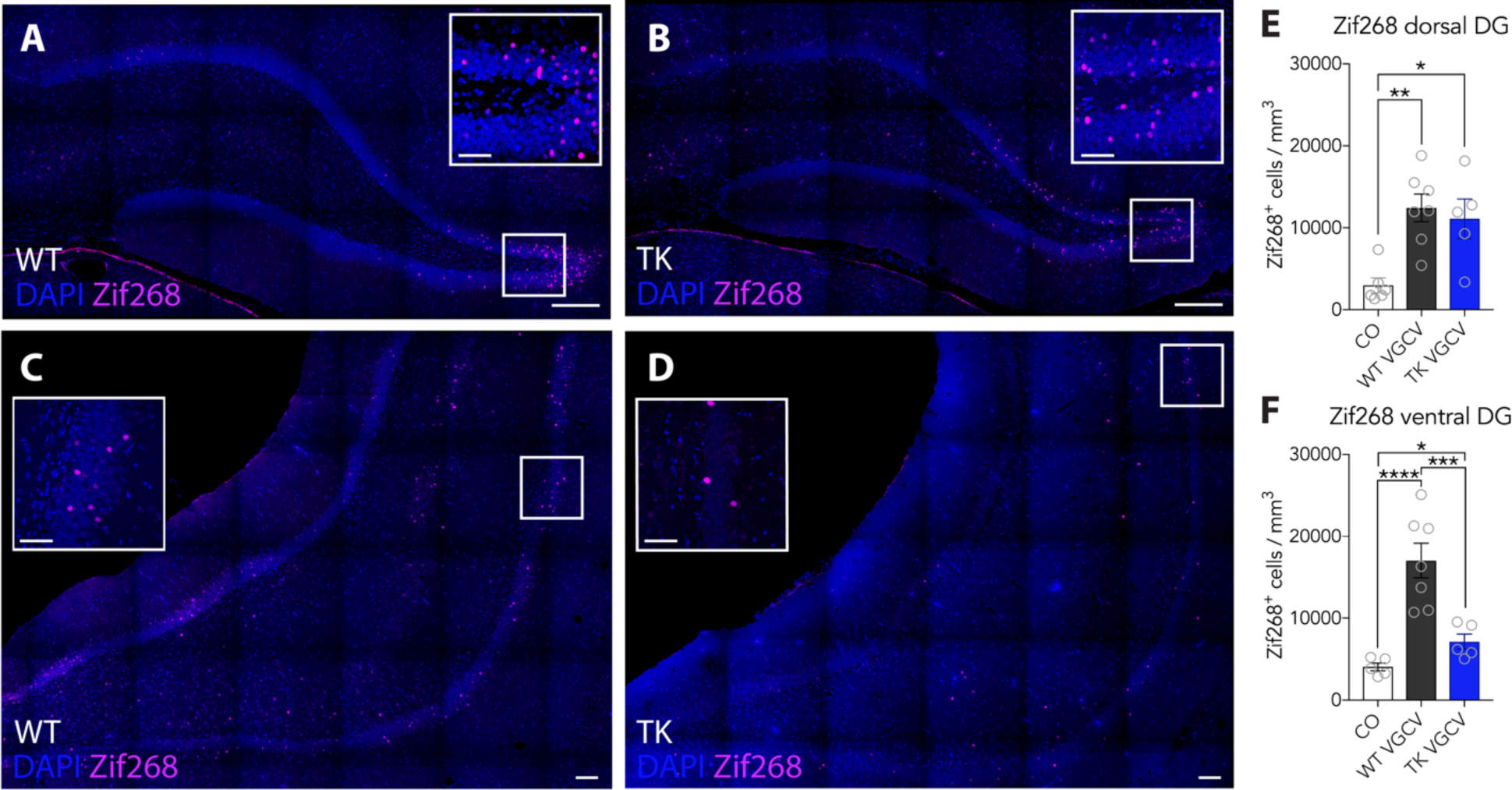
Loss of neurogenesis decreases activity in the ventral DG A-D) Representative images of the Zif268 staining in WT and TK rats in the dorsal and ventral DG 35min after the last day of performing the DD task. **E)** Performing the DD task increased expression of Zif268 in the dorsal DG in WT and TK rats compared to controls (ANOVA; effect of group: F_2,15_=9.27, P=0.0024, n=5-7; post-hoc Tukey’s test: CO 2,978 vs. WT VGCV 12,421 cells, P=0.0027; CO 2,978 vs. TK VGCV 11,112 cells, P=0.0147; WT VGCV 12,421 vs. TK VGCV 11,112 cells, P=0.8547). **F)** While WT rats had a robust (4.2x) increase in Zif268 expression in the ventral DG, the increase in TK rats was modest (1.8x) and Zif268^+^ cell densities were significantly lower than WTs (ANOVA; effect of group: F_2,14_=33.03, P<0.0001, n=5-7; post-hoc Tukey’s test: CO 4,052 vs. WT VGCV 17,017 cells, P<0.0001; CO 4,052 vs. TK VGCV 7,138 cells, P=0.0283; WT VGCV 17,017 vs. TK VGCV 7,138 cells, P=0.0008). Data are presented as mean ± standard error. Scale bars, 200µm; scale bars insert, 50µm. DG, dentate gyrus. n indicates number of rats. *, P<0.05; **, P<0.01; ***, P<0.001, ****, P<0.0001.

Adult-born neurons are sensitive to experience and undergo plastic morphological changes in response to spatial learning^23^. However, it is unknown whether decision-making processes engage new neurons and induce structural changes. We therefore labelled newborn neurons with an eGFP-expressing retrovirus and examined morphological features after rats underwent DD training, No-Delay (ND) training, or remained undisturbed in their home cage (CO) (Supplementary Fig. 5). Rats received retrovirus injections at 7 weeks prior to training, to label the oldest, most mature, cohort of neurons (“mature”) that were depleted in the TK experiment, and 3 weeks prior to DD training, to label neurons that would be young and highly-plastic at the time of training onset (“immature”).

Consistent with a role in promoting choice of delayed rewards, dendritic complexity of mature adult-born neurons was selectively increased by DD training, as measured by Sholl analyses, in both the dorsal and ventral DG (Fig. 3A-C). Neurons in DD-trained rats also had greater total dendritic length than neurons in the ND and CO groups, indicating that learning about delayed rewards specifically recruits and promotes the growth of adult-born neurons (Fig. 3D). The number of dendritic branch points was not affected by experience but was higher for all groups in the ventral compared to the dorsal DG (Supplementary Fig. 6). DD-induced dendritic plasticity was specific to mature neurons, since no effects were observed in immature adult-born neurons (Supplementary Fig. 7).

**Figure 3:**
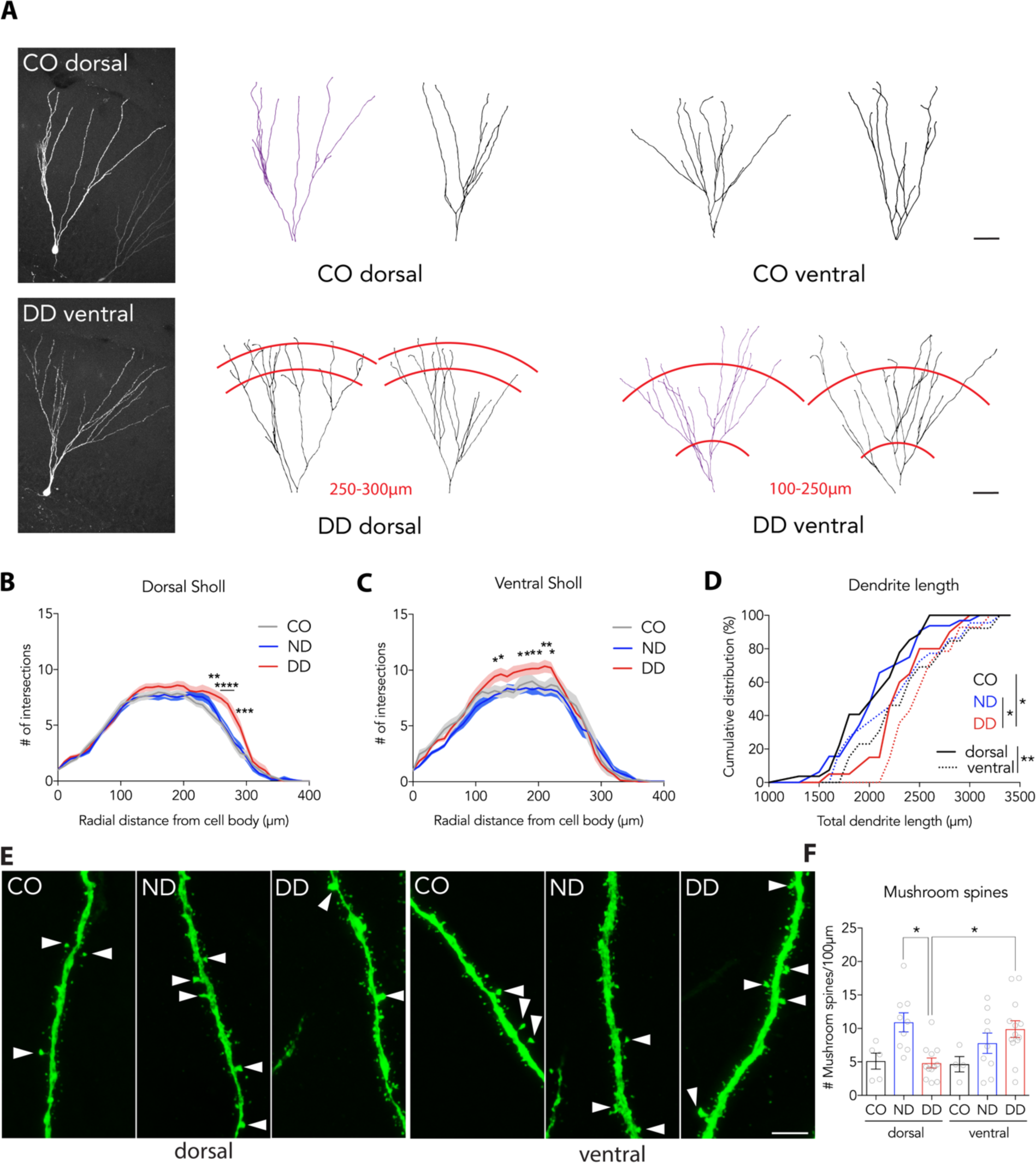
Delay-based decision-making increases dendritic complexity in new neurons and restructures spines along the dorsoventral axis. **A)** Representative confocal z-stack projections and traces of mature adult-born neurons in the dorsal and ventral DG of CO rats and rats run in the DD task. Purple traces reflect neurons in microscope images. Red lines highlight radii where differences in Sholl analyses were found between DD and both control groups. **B-C)** Sholl analysis of mature adult-born neurons revealed increased dendritic complexity in the dorsal (B) and ventral (**C**) DG from DD rats compared to CO and ND rats (dorsal: group x radial distance RM ANOVA; effect of group: F_2,75_=4.19, P=0.019; effect of radial distance: F_41,3075_=310, P<0.0001; interaction: F_82,3075_=1.965, P<0.0001; n=25-33; post-hoc Tukey’s test: ND vs. DD at 250µm, P=0.00250; at 260-280µm, P<0.0001; at 290µm, P=0.0003) (ventral: group x radial distance RM ANOVA; effect of group: F_2,46_=2.23, P=0.1196; effect of radial distance: F_41,1886_=175, P<0.0001; interaction: F_82,1886_=1.31, P=0.035; n=15-23; post-hoc Tukey’s test: ND vs. DD at 130-140µm, 170-200µm, 220µm, P<0.05; at 210µm, P<0.01). D) New neurons in DD rats had greater total dendritic length than those in CO and ND groups (group x DG region ANOVA; effect of group: F_2,122_=4.17, P=0.018; effect of DG region: F_1,122_=10.5, P=0.0015; interaction: F_2,122_=0.37, P=0.69; n=15-33; post-hoc Sidak’s test: CO 2175 vs. ND 2186µm, P=0.99; CO 2175 vs. DD 2431µm, P=0.0191; ND 2186 vs. DD 2431µm, P=0.0166). **E**) Dendritic segments from immature adult-born neurons in the dorsal and the ventral DG. Arrows indicate mushroom spines. **F**) Whereas mushroom spine density was similar in the dorsal and ventral DG in ND rats, DD training shifted mushroom spine density along the dorsoventral axis, such that dorsal neurons had fewer mushroom spines than ventral neurons, and dorsal neurons from DD rats had fewer mushroom spines than dorsal neurons from ND rats (group x DG region ANOVA; effect of group: F_2,46_=4.194, P=0.0212; effect of DG region: F_1,46_=0.1806, P=0.6728; interaction: F_2,46_=6.309, P=0.0038; n=5-13; post-hoc Tukey’s test: dorsal ND 10.92 vs. dorsal DD 4.83 mushroom spines, P=0.0090; dorsal DD 4.83 vs. ventral DD 9.9 mushroom spines, P=0.0206). Scale bars, 50µm (A) and 5µm (E). Data are presented as mean ± standard error. Significance indicated between treatment groups ND and DD in C and D. n indicates number of neurons or dendrite segments analyzed. *, P<0.05; **, P<0.01.

To determine whether DD training altered putative synaptic integration of adult-born neurons, we quantified dendritic spines in the molecular layer, where the perforant path terminates onto newborn DG neurons (Fig. 3E). We did not observe any effects of DD training on total spine density (Supplementary Fig. 8). However, DD training redistributed mushroom spines in immature neurons along the dorsoventral axis (Fig. 3F). Whereas neurons in ND-trained rats had similar numbers of mushroom spines in the dorsal and ventral DG, neurons in DD-trained rats had more mushroom spines specifically in the ventral DG than in the dorsal DG, aligning with findings that delayed reward choice is regulated by the ventral hippocampus^15^. That mushroom spines were reduced in the dorsal DG of DD-trained rats as compared to controls suggests that DD training may in fact reduce or delay the integration of new neurons in this subregion. DD-induced spine plasticity was specific to immature adult-born neurons as no differences were observed in mature adult-born neurons (Supplementary Fig. 8).

Here we identify a novel role for adult hippocampal neurogenesis in decisions about immediate vs. delayed rewards. Our findings fit with an emerging role for the human hippocampus in imagination^10^ and the mental construction of future events^9^, which may enable individuals to assign value to delayed outcomes^11^. Rodent data also indicates that the hippocampus processes information that is relevant for choices about the future: hippocampal place cell firing patterns are modulated by goals^24^ and reflect future navigational choices^5,6,25^. Notably, a population of DG neurons are selectively responsive to reward, and trigger future-oriented signals in downstream CA3^7^. Our findings suggest that new neurons might contribute a similar function and enable rats to envision the delayed consequences of pressing the high-reward lever in the DD task. Given the pervasiveness of short-sighted decision-making in psychiatric disorders^20^ our findings identify a specific behavioral function for adult neurogenesis that has broad implications for mental health.

## ACKNOWLEDGEMENTS

This work was supported by CIHR Project Grants (JSS and SBF), a NARSAD Young Investigator Grant from the Brain and Behavior Research Foundation (DRS) and a postdoctoral fellowship from the German Research Foundation (DFG) (DRS).

## METHODS

### Animals

All procedures were approved by the Animal Care Committee at the University of British Columbia and conducted in accordance with the Canadian Council on Animal Care guidelines regarding humane and ethical treatment of animals. Experimental wild type (WT) Long-Evans rats and transgenic rats expressing HSV-TK (TK) under the human GFAP promoter on a Long-Evans background were generated in the Department of Psychology animal facility with a 12-hour light/dark schedule and lights on at 9.00 am. Wild type breeders were received from Charles River, Canada. Experiments were performed during the light phase of the light/dark cycle. Breeding occurred in large polyurethane cages (47 × 37 × 21 cm) for WT and TK rats and in Opti-rat cages for WT Long-Evans rats, used for the irradiation and running experiment, containing a polycarbonate or plastic tube, aspen chip bedding and ad libitum rat chow and water. Breeders (both male and female) remained with the litters until P21, when offspring were weaned to 2 per cage in smaller polyurethane bins (48 × 27 × 20 cm) and transgenic rats were genotyped afterwards.

### Experimental procedures

#### TK rat model

In transgenic hGFAP-TK rats neurogenesis was suppressed by giving 4mg Valganciclovir (VGCV) to each rat twice per week for 6 weeks starting at 6 weeks of age. 12-week-old male TK and WT littermates were used for behavioral testing, lasting ∼5 weeks, at which point rats were perfused for histology (Dcx immunostaining). Animals were single housed and food deprived at 11 weeks of age over the course of one week to 90% of their initial weight at the start of the experiment. Animals were handled prior to operant training for a minimum of 5 min for 5 days by the experimenters.

#### Irradiation

We used irradiation as an alternative model to deplete neurogenesis. We irradiated 6-week-old male Long-Evans rats on 2 consecutive days with 5 Gray per day, resulting in a total dose of 10 Gray. We used a RadSource RS 2000 irradiator from RadSources Technologies, Inc., Suwanee, USA. Up to 4 animals’ hippocampi were simultaneously irradiated while their bodies were shielded with 3mm lead except a window above the hippocampus. Irradiated animals and all other groups in this experiment, including controls, received anesthesia on both days using 50mg/kg and 60mg/kg Pentobarbital on the first and second day, respectively. Side effects that were caused by irradiation: Grey hair appeared from 2 weeks after IR. Some animals developed a scab at the irradiation site. All 10 animals had to get their teeth trimmed, 2 animals received 1mg/kg Anafen, three times.

#### Running

Running was used to increase hippocampal neurogenesis. Running was initiated at 8 weeks of age. Animals were housed in groups of 2-3 animals in a large cage with constant access to a rat running wheel for 4 weeks. Distance animals ran was recorded daily using bicycle odometers. All animals (CO, IR, RUN) received a single dose of 5-Bromo-2’-deoxyuridine (BrdU, 200mg/kg) i.p. to label dividing neural progenitors in order to determine effectiveness of treatments at 9 weeks of age, 3 weeks after irradiation and 11 days after initiation of running, respectively. Histology was performed at the end of behavioral testing.

### Retrovirus production

The retroviral vector used in this study was derived from a Moloney Murine Leukemia-Virus (MMLV), in which eGFP expression is driven by a Ubiquitin (Ubi) promoter. Retroviral Ubi-eGFP (MMLV-eGFP; kindly provided by Dr. Shaoyu Ge) and VSV-G (kindly provided by Dr. Ana Martin-Villalba) plasmids were transfected in HEK293-GP cells (kindly provided by Dr. Diane Lagace) using PEI. Retrovirus was harvested 2 and 3 days after transfection, followed by ultracentrifugation (2 h at 27,000rpm). Retroviruses were produced as described previously ^1^. Viral titers ranged from 1 to 8 x 10^6^ colony forming units/ml.

### Stereotaxic surgery and behavioral testing

Rats were handled for 5 days the week before surgery for 3 min per day. At 6 or 10 weeks of age rats were anesthetized using isoflurane and oxygen and injected bilaterally in the dorsal (anteroposterior = − 4.0 mm; mediolateral = ±3.0 mm; dorsoventral = −3.5 mm from skull) and ventral (anteroposterior = −6.0 mm; mediolateral = ±4.7 mm; dorsoventral = −6.5 mm from skull) hippocampus using standard stereotaxic techniques. Bregma was measured at the top of the skull. 1µl of virus suspension was injected at each site at 200 nl/min. Local injection of Bupivacaine (8 mg/kg) was used as analgesic prior to surgery. Anafen (5 mg/kg) was given once a day for 3 days starting just before the surgery. Animals had at least 5 days to recover after the surgery before being food deprived for behavioral testing. Rats were food deprived at 11 weeks of age and operant training started at 12 weeks of age. Training on the ND and DD tasks was performed for 4 weeks. Brain tissue was collected 35min after animals finished with operant testing on the last day of training.

### Behavior

All animal testing was conducted in 24 operant chambers (30.5 cm x 24 cm x 21 cm; Med Associates, St Albans, VT, USA) enclosed in sound-attenuating boxes. Each box was equipped with a fan to provide ventilation and mask external noise. The chambers were equipped with two retractable levers on either site of a central food receptacle where food reinforcement (45 mg sugar pellet; Bioserv, Frenchtown, NJ, USA) was delivered by a pellet dispenser. The chambers were illuminated by a 100 mA house light located on the top center of the wall opposite the levers. Four infrared photocell sensors were positioned on the walls adjacent to the levers. Locomotor activity was indexed by the number of photo beam breaks that occurred during a session. The food receptacle contained an infrared head entry detector to determine the number of nosepokes. All experimental data were recorded by personal computers connected to chambers through an interface.

### Initial lever-press training

On the day before their first exposure to the operant chambers, each animal received ∼25 reward pellets in their home cage. On the first day of training, rats were in the operant chamber for 30 min and every 30 s 1 reward pellet was delivered into the food receptacle. On the second day of training, the food receptacle contained 2-3 reward pellets and crushed pellets were placed on the extended lever before each rat was placed in the chamber. First, rats were trained to press one of the levers to receive a reward on a fixed-ratio 1 (FR1) schedule to a criterion of 60 presses in 30 min. Levers were counterbalanced left/right between subjects. When the criterion was met, FR1 training was conducted on the other lever to ensure that both levers were experienced.

### Delay discounting pretraining

Rats were run on the following simplified version of the full task before starting the actual discounting task. These 90-trial sessions started with the levers retracted and the operant chamber in darkness. Every 30s, a new trial was initiated by the extension of one of the two levers into the chamber. If the rat failed to respond to the lever within 10s, the lever was retracted, the house light was extinguished and the trial was scored as an omission. A response within 10s of lever insertion resulted in delivery of a single pellet. In every pair of trials, the left or right lever was presented once, and the order within the pair of trials was random. Rats were trained for 3-5 days on this task to a criterion of 80 or more successful trials (i.e. ≦10 omissions).

### Reward Magnitude Discrimination/No Delay task

The No Delay task (ND) is a full version of the delay discounting task that the animals were trained on, except that the low (1 pellet) and the high reward (4 pellets) were both delivered immediately after 1 lever press. Animals from the Delay Discounting experiments were trained on this task for 2 consecutive days before the actual task. Left and right levers were counterbalanced between groups for being the high reward lever. Each day, animals received one training session with 4 blocks of 12 trials. Each block consisted of 2 forced choice trials (only one lever extends) and 10 free choice trials (both levers extend). Every 70s the house light came on and one or both levers extended. If the rat didn’t press a lever during the next 10s, levers retract and it was counted as an omission. Animals merely had to choose between a low and a high reward and thus, within 2 days, quickly learned to prefer the high reward over the low reward.

Animals in the ND group of the virus experiment continued to be trained on this task for 28 days.

### Delay discounting

Subsequently, after the 2-day Reward Magnitude Discrimination/No Delay task, animals received daily training sessions on the Delay Discounting (DD) task. Like in the ND task, one session consisted of 48 trials, divided into 4 blocks ^2^. The lever (left or right, counterbalanced between groups) that was assigned to the high reward (HR) in the ND task remained associated with the HR in the DD task. One block started with 2 forced choice trials, where only one of the reward levers was extended (one trial for each lever, presented randomly), followed by 10 free choice trials. The inter trial interval (ITI) was 70s, regardless of lever choice, leading to a total of 56min per session. At the beginning of each choice trial, the house light was illuminated and both levers extended after 2 s. If the rat failed to respond within 10 s, similar to the lever-press training, both levers would retract, the trial would be scored as an omission, houselights would go off until the next scheduled trial would begin. On each choice trial, a press on the low reward (LR) lever retracted both levers and delivered one sugar pellet immediately. Choice of the HR lever also retracted both levers immediately, but 4 pellets were delivered after a delay, which increased over the 4 blocks of trials (0 s, 15 s, 30 s, 45 s). During the delay, the chamber remained in darkness as in the ITI and was re-illuminated during the reward delivery at the end of the delay. Pellets were delivered 0.5 s apart. After the delivery of the reward, the house light remained lit for another 4 s before it returned to ITI state. Daily training sessions continued for 7 days a week until behavior stabilized, which was achieved at about 28 to 31 days. If expression of immediate early genes (IEGs) or morphology of virally labeled neurons was assessed in brain tissue, on the last day of the experiment, animals were perfused and brain tissue extracted 35 min after animals finished the DD task.

### Restraint stress

Acute stress was induced by restraining rats for 30 min in a Plexiglas Cylindrical Restrainer tube (Broome style rodent restrainer 250 g to 500 g; 6.35 cm x 6.35 mm x 21.6cm; Plas Labs, Inc.^™^), in a quiet, lit, and ventilated room ^3^. After rats were placed in the tubes, restrainer length was adjusted to keep rats immobilized without causing pain. Upon being released from the tube into their home cage, rats were immediately transported to the operant chamber to start the DD task.

### Immunofluorescence

Animals were anesthetized using isoflurane and transcardially perfused with 4 % paraformaldehyde (PFA) in PBS. Brains were extracted and post-fixed for 48 h in 4 % PFA. To analyze IEG activity or assess levels of neurogenesis, tissue was cut at 50 µm (series of 10) on a vibratome. To examine neuronal morphology of virally labeled new-born neurons, tissue was cut at 100 µm (series of 8) on a vibratome. Cut tissue was stored in anti-freeze at −20 °C until further processing.

Free floating sections were washed three times in PBS. When staining for BrdU, we additionally incubated sections in 2N HCl for 30min at RT and washed tissue 5 times in PBS for 3min. Then, for all stainings, sections were incubated in blocking solution (3 % horse serum and 0.5 % Triton-X). Primary antibodies mouse anti-GFP (DSHB, GFP-12E6, 1:100), rabbit anti-Zif268 (Santa Cruz, sc189, egr-1, 1:1000), goat anti-Dcx (Santa Cruz, C-18, 1:200), goat anti-TK (Santa Cruz, sc28038, 1:200), mouse anti-BrdU (BD Biosciences, 347580, 1:200) were incubated at 4 °C for 72 h. Sections were washed three times and then incubated with secondary antibodies (1:400, Invitrogen) in blocking solution for 2 h at 4 °C (donkey anti-mouse Alexa488, donkey anti-goat Alexa555, donkey anti rabbit Alexa647). Subsequently, sections were washed once, nuclei were stained with DAPI (1mg/ml) 1:1000 in PBS for 5 min at room temperature and washed in PBS another 3 x for 5 min. Sections were mounted on glass cover slides and cover slipped with PVA-Dabco to preserve fluorescence.

### Cell counting

Confocal images of the dorsal and ventral dentate gyrus were taken using a water 25 x objective on a Leica SP8 microscope. Settings were 1024×1024 pixels, zoom 1×, imaging speed was 200Hz, and z height was 1µm. Analysis of Dcx and BrdU positive cells in the GCL layer of the dorsal and ventral hippocampal DG were performed using the Cell Counter Plugin from ImageJ. Cells densities were calculated by dividing the number of positive cells in the GCL of the DG by the GCL volume.

### Neuronal morphology analysis

Confocal images of new-born neurons in the dorsal and ventral dentate gyrus were taken using a water 25 x objective on a Leica SP8. Settings were 1024×1024 pixels, zoom 1×, imaging speed was 200Hz, and z height was 1µm. Simple Neurite Tracer (Image J) was used to trace dendritic trees of newborn granule cells to collect data for Sholl analysis and total dendritic length. Branch points were manually counted from the traces by the experimenter. Five neurons were recorded per GFP positive animal in the dorsal and ventral hippocampus. Neurons with more than 3 primary and secondary dendrites cut due to tissue processing and microscope imaging were excluded from the analysis.

### Spine analysis

Confocal images of spines from newly generated virus labeled neurons were taken using a glycerol 63 x objective on a Leica SP8 with 5 x zoom and 1µm z steps. Settings were 1024×1024 pixels, zoom 5×, imaging speed was 100Hz, and line average was 2. 5 dendrite segments from different neurons were recorded per GFP positive animal in the dorsal and ventral hippocampus. Average dendrite length was (40 µm). Dendrite length was measured using Simple Neurite Tracer (Image J). Spines were counted manually by a human experimenter blind to the experimental conditions using the Cell Counter plugin in ImageJ. Mushroom spines were measured with ImageJ and defined as having a diameter ≧0.6 µm.

### Zif268 analysis

Confocal images of the hippocampal DG area as well as images of the Nucleus Accumbens (NAc) were taken using a water 25 x objective on a Leica SP8. Settings were 1024×1024 pixels, zoom 1×, imaging speed was 200Hz, and z height was 1µm. Intensities of Zif268 staining in labeled cells in the Core and the Shell of the NAc were analyzed measuring the mean gray value using ImageJ. Counts of single Zif268 positive cells being at least 4x as bright as the background in the hippocampal DG were done manually by an experimenter blind to the experimental conditions using ImageJ. Cells densities were calculated dividing the number of Zif268 positive cells in the supra-or infrapyramidal blade and dividing it by the respective volume.

### Statistical Analyses

All data were analyzed using Graph Pad Prism 7 software. We used unpaired T-tests and one-way or two-way ANOVAs to analyze effects of genotypes and treatments on neurogenesis, IEG expression, neuronal morphology and behavior. When we observed significant interactions, we used Sidak’s, Tukey’s or Dunnett’s tests to compare groups as indicated. If distributions failed tests of normality and homogeneity of variance, analyses were run on log transformed data (but graphs display untransformed data). In all cases, statistical significance was set at P=0.05.

### Data Availability

The datasets generated during and/or analysed during the current study are available from the corresponding author on reasonable request

## SUPPLEMENTARY FIGURES

**Supplementary Figure 1:**
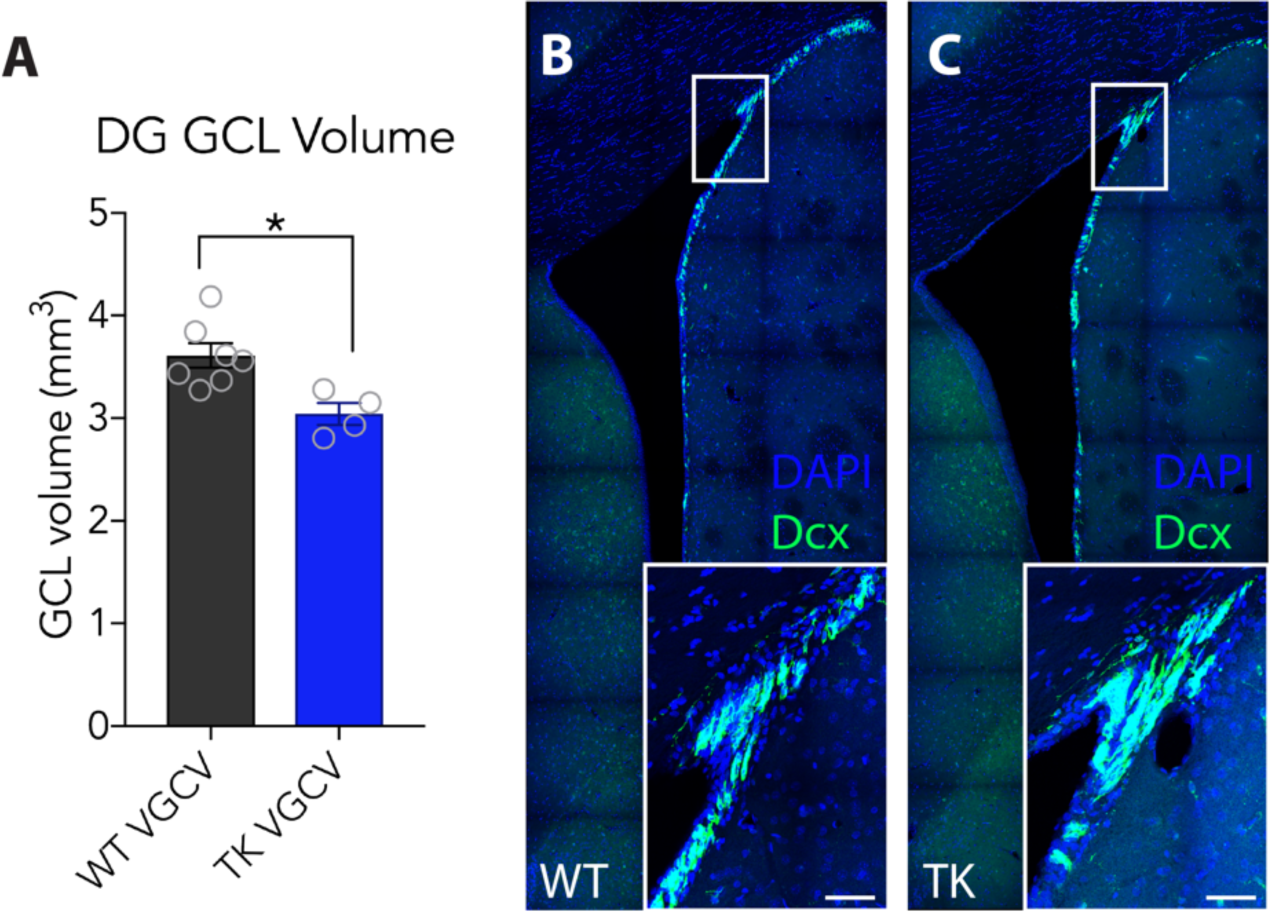
Histology in VGCV-treated TK rats. **A)** VGCV treatment led to a significant reduction in the volume if the DG granule cell layer (T-test; WT 3.6 vs. TK 3.0 mm^3^, P=0.0109, n=4-7). **B, C)** Immature, DCX^+^ neurons appeared normal in the subventricular zone. Data presented as mean ± standard error. Scale bars, 50µm. DG, dentate gyrus; GCL, granule cell layer. *, P<0.05.

**Supplementary Figure 2:**
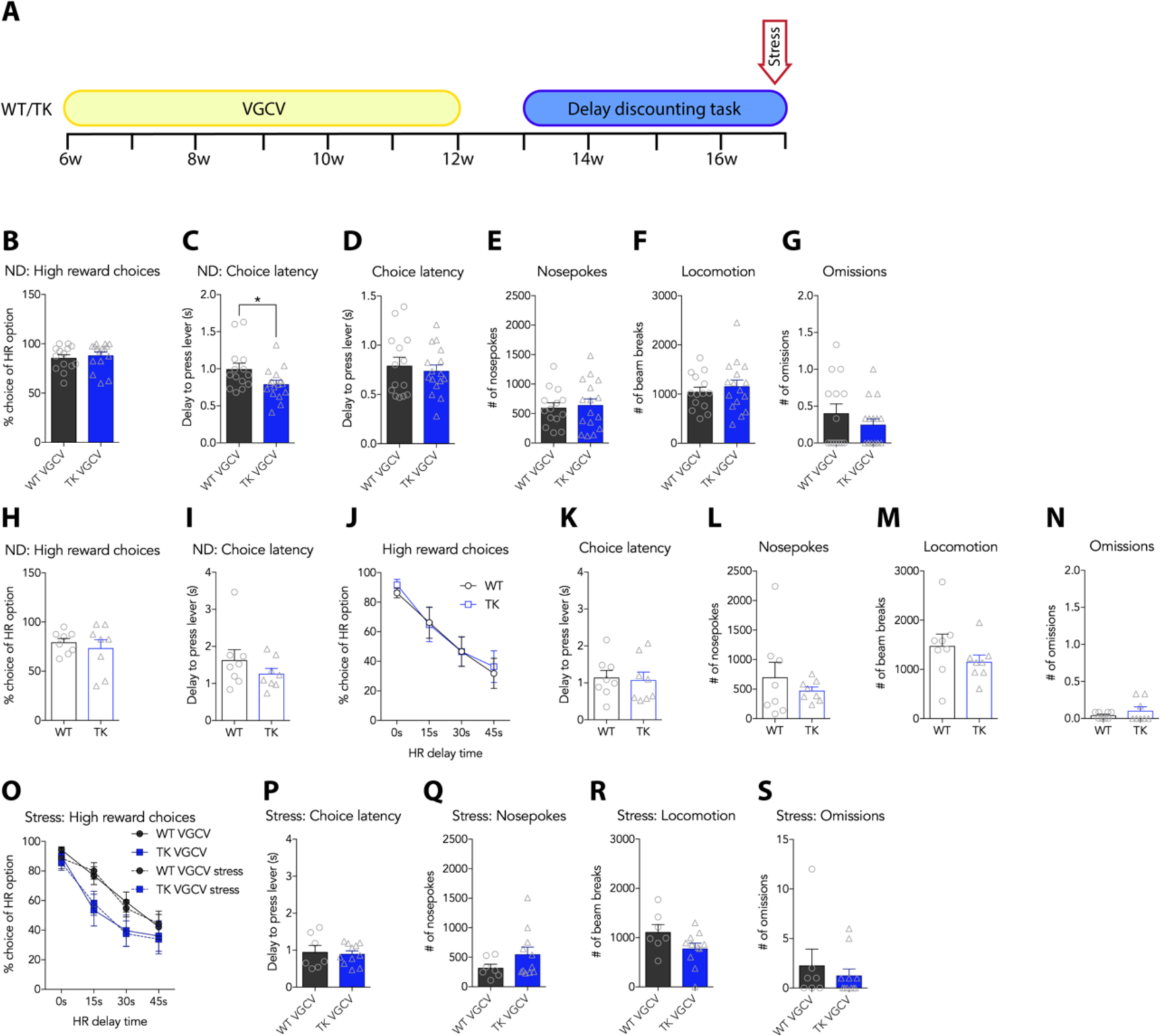
Delay discounting task behaviors in VGCV-treated, stressed VGCV-treated & untreated rats. **A)** Timeline of VGCV treatment and testing in the DD task. **B-G)** Additional behavioral analyses of VGCV-treated WT and TK rats. B) WT and TK rats did not differ in preference for high reward in a No Delay (ND) pretraining version of the DD task (T-test; P=0.57, n=14-16). C) Choice latency was slightly lower in TK rats in the initial reward magnitude discrimination task/no delay version of the DD task (T-test; WT 0.996 vs. TK 0.793s, P=0.037, n=14-16). **D-G)** The latency to press the lever (T-test; P=0.61, n=14-16), the number of nosepokes (T-test; P=0.73, n=14-16), locomotion (T-test; P=0.49, n=14-16), and the number of omitted trials (T-test; P=0.29, n=14-16) were not affected by lack of neurogenesis. **H-N)** DD task behaviors in WT and TK rats that did not receive VGCV (i.e. neurogenesis-intact). H) Untreated WT and TK rats showed similar preference for a high reward in a no delay version of the DD task (T-test; P=0.55, n=8). I) There was no difference in choice latency between untreated WT and TK rats (T-test; P=0.27, n=8). J) Delayed reward preferences were not different between untreated WT and TK rats (genotype x time RM ANOVA; effect of genotype: F_1,14_=0.036, P=0.85; effect of time: F_3,42_=39.98, P<0.0001; interaction: F_3,42_=0.20, P=0.90; n=8). **K-N)** The latency to press the lever (T-test; P=0.82, n=8), the number of nosepokes (T-test; P=0.41, n=8), locomotion (T-test; P=0.27, n=8), and the number of omitted trials (T-test; P=0.29, n=8) were not different between untreated WT and TK rats. **O-S)** 30 min restraint stress did not alter VGCV-treated WT or TK behavior in the DD task. Data show the day of stress compared to a normal test day prior to stress. O) DD in WT and TK rats (treatment x time RM ANOVA; WT vs. WT stress, effect of stress, P=0.73; TK vs. TK stress, effect of stress, P=0.79). **P-S)** Choice latency (T-test; P=0.76, n=8), the number of nosepokes (T-test; P=0.18, n=8), locomotion (T-test; P=0.083, n=8), and the number of omissions (T-test; P=0.52, n=8) were not affected by 30 min restraint stress in VGCV-treated TK or WT rats. Data shown represent the average of the last 3 test days. For ND training, data from single last training day is shown. Data are presented as mean ± standard error. HR, high reward; ND, no delay version of the DD task. *, P<0.05.

**Supplementary Figure 3:**
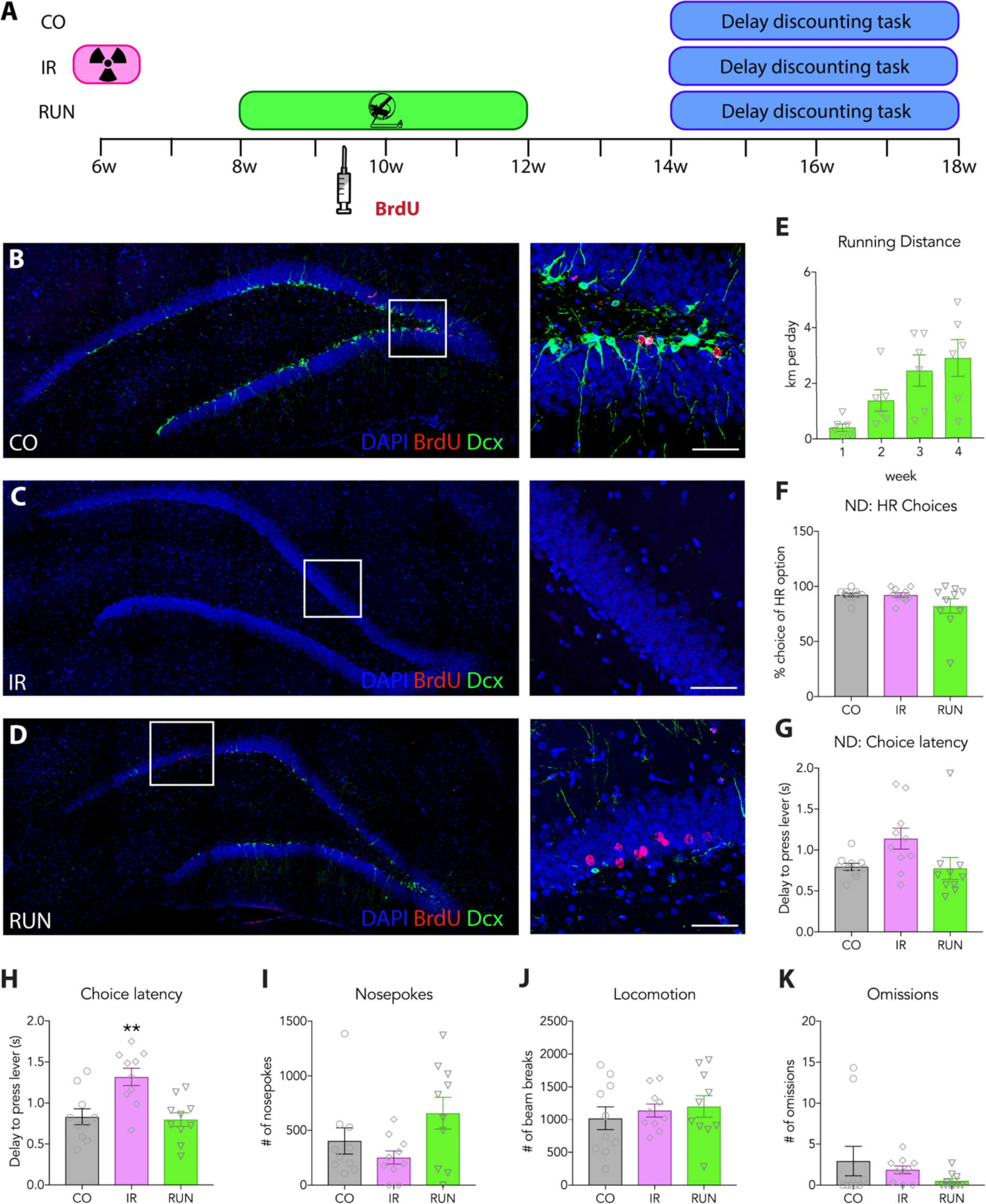
Effects of running and irradiation on neurogenesis and additional behaviors in the delay discounting task. **A)** Schematic overview of the experimental timeline. **B-D)** Representative images of BrdU-labeled neurons and Dcx^+^ immature neurons in control (CO), irradiated (IR) and running (RUN) animals. **E)** Running distance increased over the 4 weeks of continuous wheel access (RM ANOVA with Geisser-Greenhouse’s correction; effect of time: F_1.4,6.9_=13.04, P=0.0066, n=6 cages). **F)** Irradiation (IR) or running (RUN) did not affect the general preference for the high reward in a no-delay version of the DD task (ANOVA; effect of group: F_2,27_=2.00, P=0.15, n=10). **G)** Choice latency was not significantly affected in IR or RUN rats compared to controls (CO) in a No Delay version of the DD task (ANOVA; effect of group: F_2,27_=3.46, P=0.046, n=10). **H)** Choice latency was greater in IR rats compared to controls in the DD task (ANOVA; effect of group: F_2,27_=9.2, P=0.0009, n=10). **I-K)** The number of nosepokes (ANOVA; effect of group: F_2,27_=3.21, P=0.056, n=10), locomotion (ANOVA; effect of group: F_2,27_=3766, P=0.69, n=10) and number of omitted trials (ANOVA; effect of group: F_2,27_=1.23, P=0.31, n=10) were not affected in IR or RUN rats compared to controls. Data shown represent the average of the last 3 test days. For No-Delay training, data from single last training day is shown. Data are presented as mean ± standard error. Scale bars, 50µm. ND, No-Delay version of the DD task; HR, high reward. **, P<0.01; ***, P<0.0001.

**Supplementary Figure 4:**
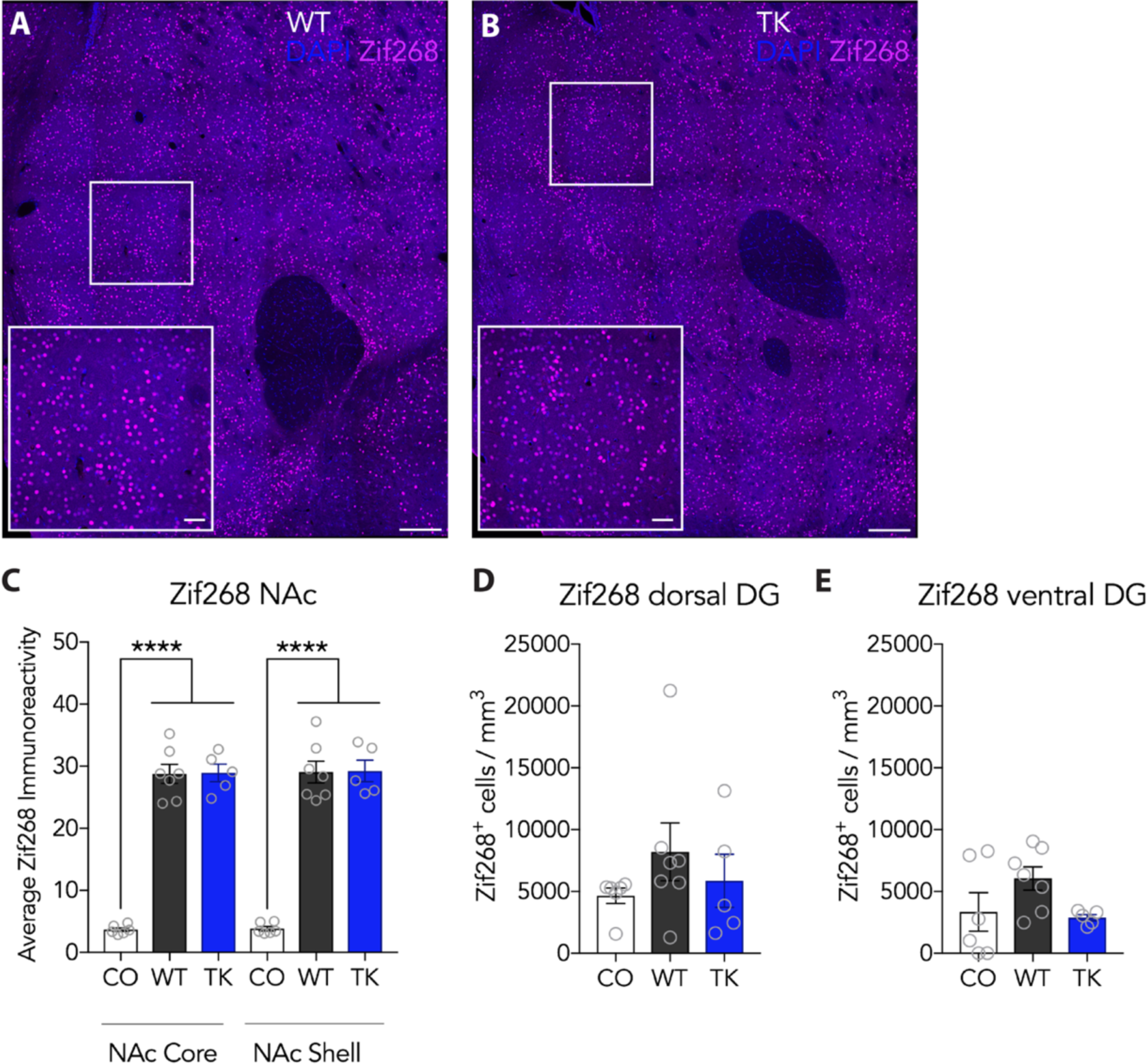
Immediate-erly gene expression after the DD task. **A-B)** Representative images of Zif268 IEG expression in the Nucleus Accumbens (NAc) in VGCV-treated WT and TK rats after performing the DD task. **C)** Zif268 expression in the Core and the Shell of NAc is significantly increased in VGCV-treated WT and TK rats after animals performed the DD task. The average intensity of the Zif268 immunoreactivity was comparable between WT and TKs (group x NAc region ANOVA; effect of group: F_2,15_=117.6, P<0.0001; effect of NAc region: F_1,15_=0.92, P=0.35; interaction: F_2,15_=0.0297, P=0.97; n=5-7). D-E) Zif268^+^ cells in the infrapyramidal blade of the dorsal (ANOVA; effect of group: F_2,15_=0.96, P=0.41, n=5-7) and ventral (ANOVA; effect of group: F_2,15_=2.49, P=0.12, n=5-7) DG were not significantly increased in WT and TK rats after performing the DD task compared to food restricted cage control animals. Scale bars, 200µm; inset scale bar, 50µm. Data are presented as mean ± standard error. ****, P<0.0001.

**Supplementary Figure 5:**
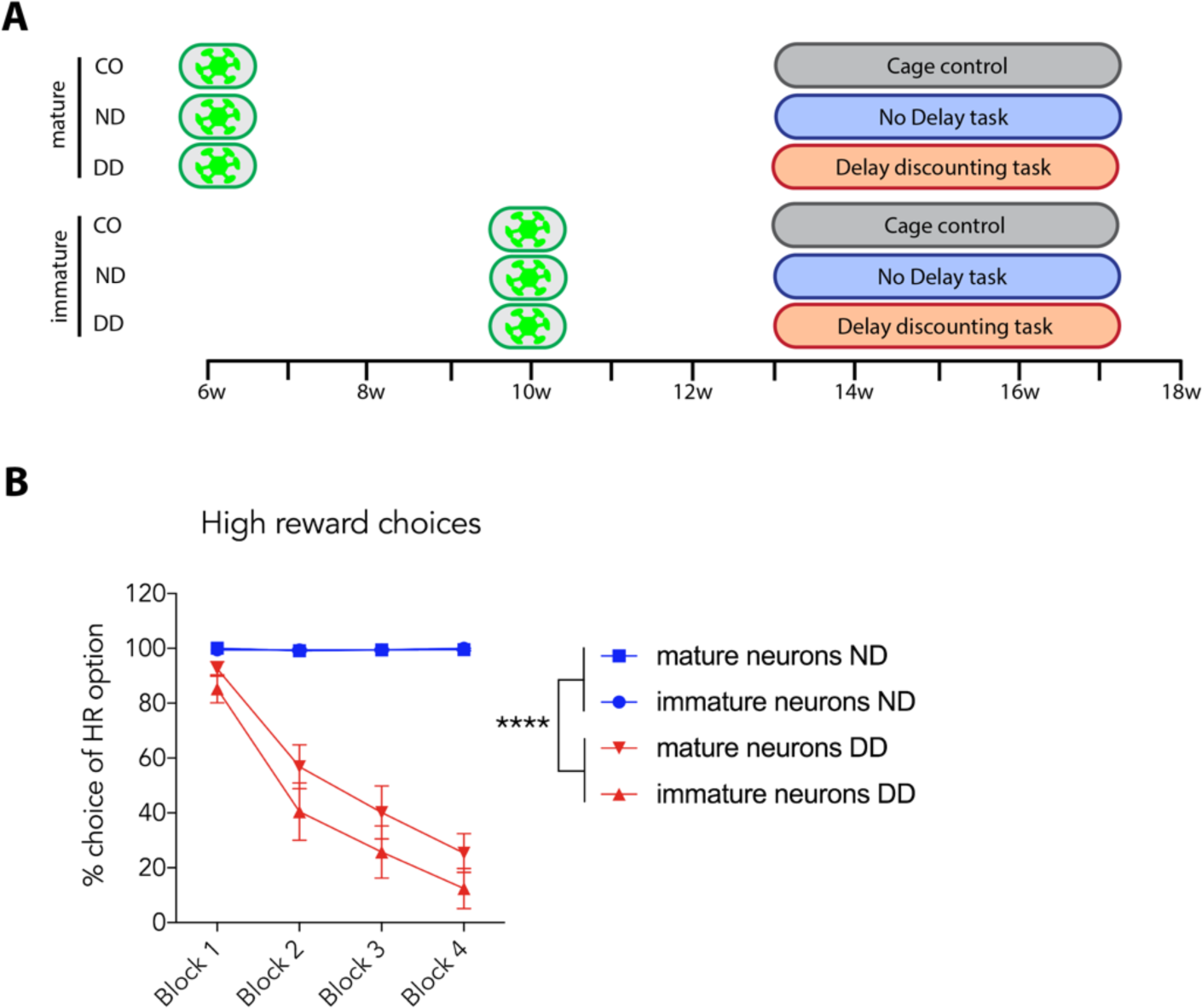
Experimental timeline and behavioral data of retrovirus-injected animals in the DD task. **A**) Experimental overview. Rats were injected with retrovirus at 7 weeks or 3 weeks before training to label cohorts of adult-born neurons that were mature or immature at the time of training onset, respectively. Virus-injected rats either remained in their home cages (CO), were trained in a no-delay version of the task (ND) or were trained in the standard delay discounting (DD) task. **B)** Rats run in the ND task showed a close to 100% preference for the high reward in all blocks, whereas rats run in the DD task showed a decrease in the preference for the high reward with increasing delay times (group x block RM ANOVA; effect of group: F_3,25_=46.5, P<0.0001; effect of block: F_3,75_=52.12, P<0.0001; interaction: F_9,75_=17.58, P<0.0001; n=6-8). Data shown represent the average of the last 3 test days. Data are presented as mean ± standard error. ****, P<0.0001.

**Supplementary Figure 6:**
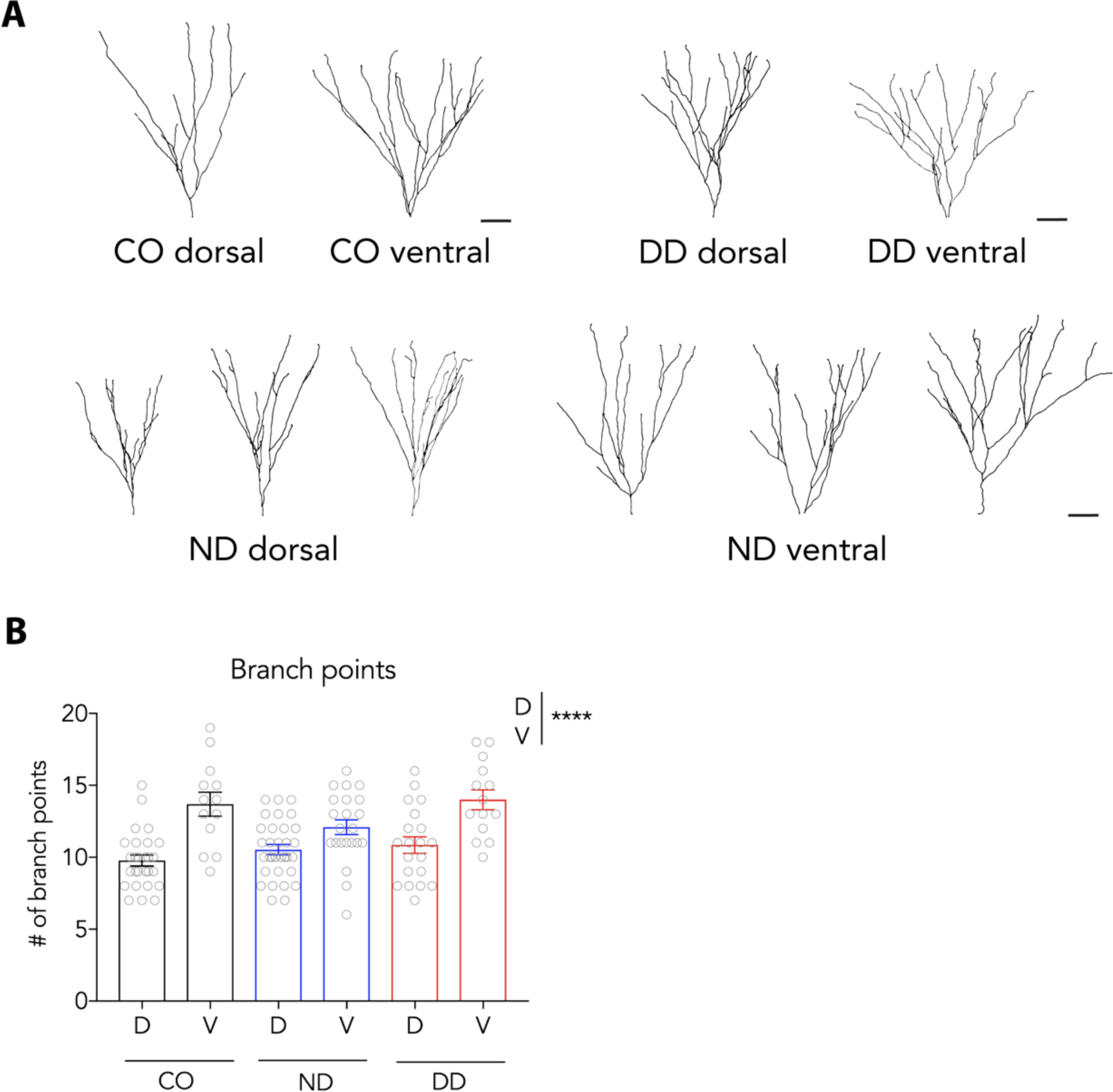
Additional morphological analyses of mature adult-born neurons. **A**) Representative traces of mature adult-born neurons in the dorsal and ventral DG of CO rats and rats run in the ND and DD tasks. **B**) There were no differences in branch points between the CO, ND and DD groups but ventral DG neurons had significantly more branch points compared to dorsal DG neurons (group x DG region ANOVA; effect of group: F_2,122_=2.27, P=0.108, effect of DG region: F_1,122_=43.15, P<0.0001; interaction: F_2,122_=2.86, P=0.061; n=15-33). Scale bars 50 µm. Data are presented as mean ± standard error. ****, P<0.0001.

**Supplementary Figure 7:**
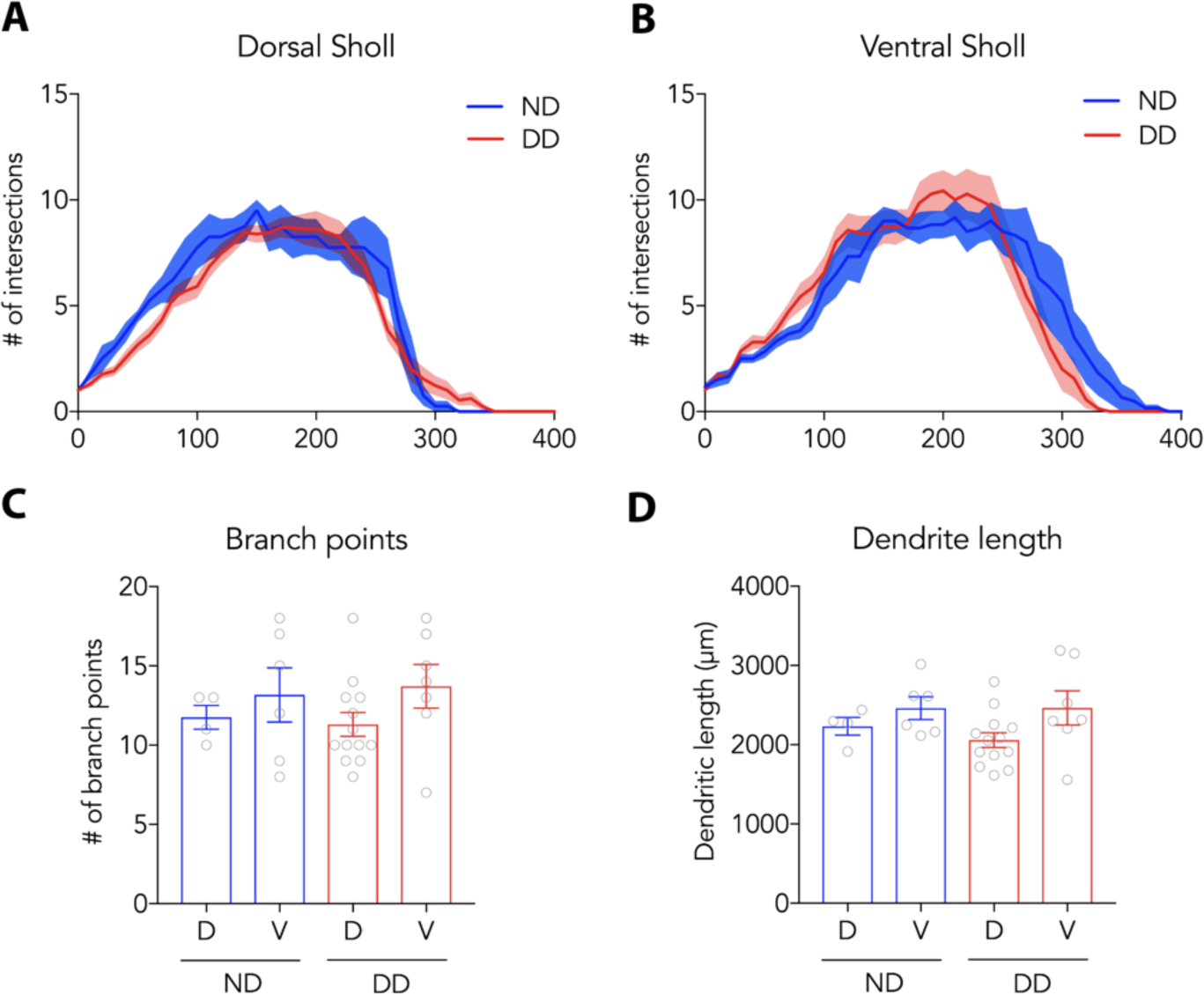
Dendritic morphology of immature neurons. **A-B)** Integration of immature neurons in the dorsal and ventral hippocampus was not affected by learning the DD task, as measured by Sholl analyses (dorsal: group x radial distance RM ANOVA; effect of group: F_1,15_=1.27, P=0.28; effect of radial distance: F_40,600_=56.86, P<0.0001; interaction: F_40,600_=0.92, P=0.61; n=4-13) (ventral: group x radial distance RM ANOVA; effect of group: F_1,11_=0.004, P=0.95; effect of radial distance: F_40,440_=42.1, P<0.0001; interaction: F_40,440_=1.476, P=0.034; n=6-7). **C-D**) Total dendritic length (group x DG region ANOVA; effect of group: F_1,26_=0.29, P=0.59; effect of DG region: F_1,26_=4.06, P=0.054; interaction: F_1,26_=0.31, P=0.58; n=4-13) and branch points (group x DG region ANOVA; effect of group: F_1,26_=0.002, P=0.97; effect of DG region: F_1,26_=2.30, P=0.14; interaction: F_1,26_=0.15, P=0.70, n=4-13) were similar between the two groups. Data are presented as mean ± standard error.

**Supplementary Figure 8:**
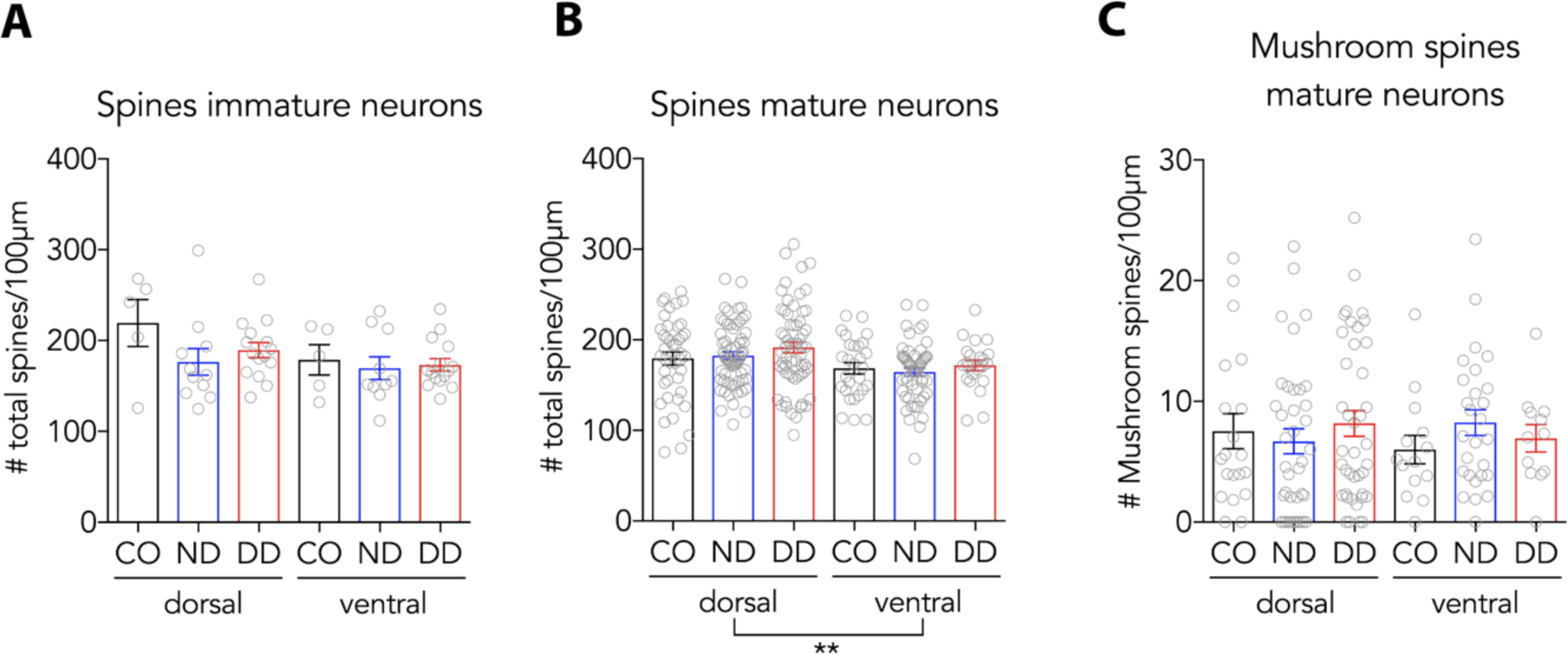
Additional analyses of spines on immature and mature adult-born neurons. **A)** Learning the DD task did not alter overall spine density in immature neurons (group x DG region ANOVA; effect of group: F_1,47_=0.65, P=0.42; effect of DG region: F_1,47_=1.23, P=0.27; interaction: F_1,47_=0.21, P=0.65; n=10-15). **B)** Mature adult-born neurons have more spines in the dorsal DG than in the ventral DG (group x DG region ANOVA; effect of group: F_2,265_=1.03, P=0.36; effect of DG region: F_1,265_=10.16, P=0.0016; interaction: F_2,265_=0.26, P=0.77; n=22-64). **C)** Operant training did not alter the number of mushroom spines compared to control animals (group x DG region ANOVA; effect of group: F_2,144_=0.19, P=0.83; effect of DG region: F_1,144_=0.13, P=0.72; interaction: F_2,144_=0.98, P=0.38; n=12-40).

